# Positive-Unlabeled Learning for Predicting Small Molecule MS2 Identifiability from MS1 Context and Acquisition Parameters

**DOI:** 10.64898/2026.01.23.701093

**Authors:** Madina Bekbergenova, Tao Jiang, Louis-Félix Nothias, Wout Bittremieux

## Abstract

**Motivation:** The quality of tandem mass spectra critically determines metabolite identifiability in untargeted metabolomics, yet optimizing MS2 acquisition parameters experimentally is costly, time-consuming, and infeasible across the full diversity of samples and instruments. A key obstacle to computational quality assessment is the absence of reliable negative labels: most MS2 spectra remain unannotated not because they are low quality, but because the corresponding compounds are absent from reference libraries. This label ambiguity fundamentally limits supervised learning approaches and complicates acquisition-time decision making.

**Results:** We present a deep learning framework that predicts the probability that an MS2 scan will be identifiable using only the preceding MS1 spectrum and instrument acquisition parameters, without inspecting the MS2 spectrum itself. The problem is formulated as positive-unlabeled learning and addressed using a non-negative positive-unlabeled objective, enabling robust training despite missing negative labels. Trained on over eight million MS2 scans from public Orbitrap metabolomics datasets and evaluated on laboratory-disjoint benchmarks, the model recovered 90% of known identifiable spectra in a held-out test set. Predicted probabilities generalized to unseen laboratories and stratified unlabeled spectra in a physicochemically meaningful manner. Independent validation demonstrated that high predicted quality is associated with increased structural explainability, richer fragmentation patterns, canonical precursor charge states, and reduced spectral interference. These results indicate that MS2 identifiability can be anticipated from precursor context and acquisition settings alone.

**Availability and implementation:** Code is available at https://github.com/bittremieuxlab/pu_ms2_identifiability. Model weights and processed datasets are available at 10.5281/zenodo.18266932.

## 1 Introduction

Liquid chromatography coupled with tandem mass spectrometry (LC-MS/MS) is the cornerstone of untargeted metabolomics, enabling large-scale detection and comparative analysis of small molecules in complex biological and environmental samples [2]. Despite continued advances in instrumentation and data analysis, confident identification remains a major bottleneck: in many untargeted studies, only a small fraction of acquired MS2 spectra can be annotated against reference libraries [5]. This limited interpretability constrains biological insight and reduces the robustness of downstream analyses such as molecular networking, pathway enrichment, and comparative metabolomics. While the incompleteness of spectral libraries is a well-recognized contributor to low annotation rates [6], the quality and informativeness of the acquired MS2 spectra themselves play an equally critical role in determining whether a compound can be identified.

In practice, MS2 “quality” is a multifaceted concept. Informative fragmentation spectra typically exhibit sufficient fragment ion richness, chemically coherent fragmentation patterns, low levels of spectral interference, and precursor isolation purity consistent with the assumed charge and adduct state. These properties are shaped by a complex interplay between precursor characteristics observable in the MS1 scan, such as intensity, local spectral congestion, and isotope or adduct patterns; and the instrument method parameters used during fragmentation, including isolation width, automatic gain control (AGC), ion injection time, resolving power, and collision energy. Suboptimal choices can result in sparse spectra, excessive noise, chimeric fragmentation, or over-fragmentation, all of which reduce the likelihood that a spectrum will support reliable structural inference or library matching.

The impact of acquisition parameters on MS2 quality has been systematically explored in controlled experimental studies, particularly for Orbitrap-based instruments, demonstrating that settings such as AGC target, resolution, and isolation width substantially affect spectral quality and annotation performance [21]. However, experimentally optimizing these parameters is expensive and time-consuming, and such studies necessarily explore only a limited subset of the parameter space, compound classes, and sample types. Given the high dimensionality of acquisition settings and their nonlinear interactions with sample complexity and precursor context, it is infeasible to exhaustively characterize optimal conditions through empirical experimentation alone. This has motivated interest in computational approaches that can guide acquisition design and parameter selection [7].

In proteomics, predictive modeling of MS2 spectra and their quality has achieved notable success [12]. Because in this setting the chemical space is comparatively constrained and identification pipelines provide clear ground truth through database search with controlled false discovery rates [10], supervised learning approaches have been proposed to predict spectrum quality or suitability for identification [13]. Importantly, the availability of large datasets with reliable negative examples has made it feasible to train and benchmark such predictors using fully supervised learning [20].

In contrast, predicting MS2 quality in untargeted metabolomics presents fundamentally different challenges. The chemical space of small molecules is vast and heterogeneous, fragmentation pathways are highly compound- and adduct-specific, and reference libraries cover only a small fraction of the metabolites encountered in real samples. Consequently, the absence of a library match does not necessarily imply poor spectral quality: high-quality, chemically informative spectra frequently remain unannotated simply because the corresponding compound is absent from existing libraries [6]. This creates a critical obstacle for supervised learning approaches, as reliable labels are available only for a subset of high-quality spectra, while the remaining data form a heterogeneous mixture of truly low-quality scans and high-quality but unidentified spectra. Treating all unmatched spectra as negatives would therefore introduce substantial label noise and bias.

Existing approaches to spectral quality assessment in mass spectrometry have largely focused on post-acquisition analysis, where the MS2 spectrum itself is used to derive quality scores or filtering heuristics [12]. While valuable for downstream processing and quality control, such methods cannot inform acquisition-time decisions or guide the selection of instrument parameters before data are collected. In metabolomics, most machine learning efforts have instead focused on improving identification given an acquired spectrum [8], through spectral similarity learning, molecular networking, or structure elucidation, rather than predicting whether a particular acquisition setting is likely to yield an informative MS2 spectrum in the first place. As a result, there remains a gap between experimental parameter optimization and computational modeling of spectral identifiability in realistic, unlabeled metabolomics datasets.

Here, we address this gap by framing MS2 identifiability prediction as a positive-unlabeled (PU) learning problem and by shifting the focus from post hoc spectral evaluation to pre-acquisition forecasting (Figure 1a). Rather than analyzing the MS2 spectrum itself, we predict the probability that an MS2 scan will be of sufficient quality to support confident identification based solely on the immediately preceding MS1 spectrum and the instrument parameters used to generate the MS2 scan. This formulation reflects the causal process underlying MS2 generation and enables prediction before or independently of MS2 interpretation. To cope with the absence of reliable negative labels, we employ a non-negative positive-unlabeled (nnPU) learning strategy [16], allowing the model to learn from confidently identified spectra while accounting for the ambiguous nature of unlabeled data. By combining large-scale public metabolomics data, instrument metadata, and a principled learning framework, this approach aims to provide a model-driven foundation for computational guidance of MS2 acquisition strategies in untargeted metabolomics.

**Figure 1.**
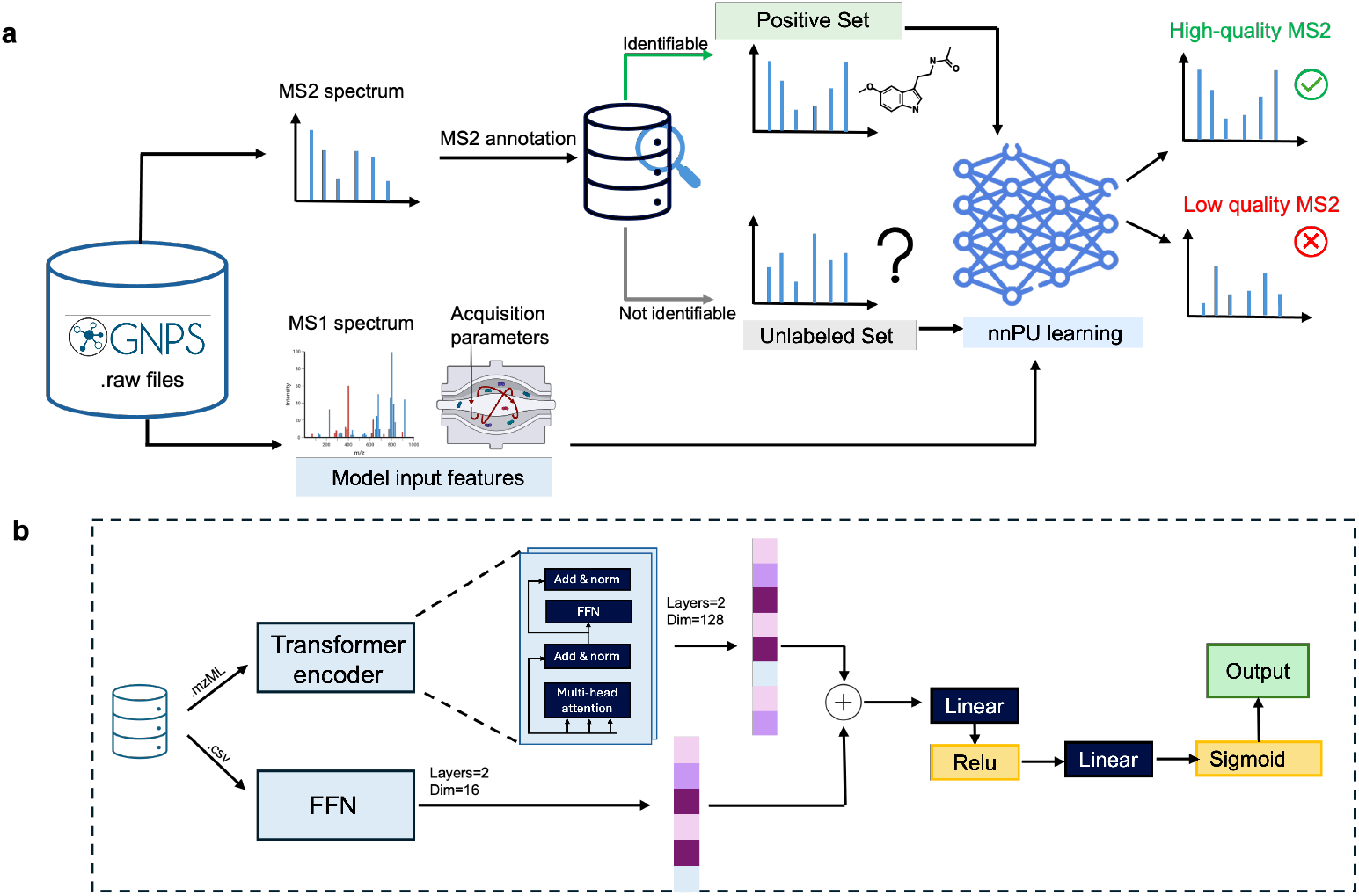
**(a)** Overall workflow of the proposed framework. For each MS2 acquisition event, the model receives as input its preceding MS1 spectrum together with the acquisition parameters used to generate the MS2 scan. The model predicts the probability that the resulting MS2 spectrum will be of sufficient quality to support confident identification. Training labels are derived from spectral library matching: MS2 scans that match reference spectra form the positive set, while unmatched scans constitute the unlabeled set. Because unmatched scans may include both low-quality spectra and high-quality spectra corresponding to compounds absent from reference libraries, the unlabeled set is treated as a mixture of positive and negative examples. The model is trained using an nnPU learning objective to account for the absence of reliable negative labels and to recover high-quality spectra hidden within the unlabeled set. **(b)** Model architecture. Preprocessed MS1 spectra, represented by *m/z* and intensity values extracted from mzML files, are encoded using a spectrum transformer-based encoder to produce an MS1 embedding. In parallel, per-scan instrument and acquisition parameters are encoded using a fully connected neural network to produce an instrument-setting embedding. The MS1 and instrument embeddings are concatenated and passed through an integration layer followed by a classification head, which outputs a scalar probability corresponding to the predicted likelihood that the MS2 spectrum generated under the given conditions will be of high quality and identifiable.

## 2 Methods

### Data sources and preprocessing

Publicly available untargeted metabolomics datasets were obtained from the GNPS/MassIVE repository [22]. Only datasets acquired on Thermo Fisher Scientific Orbitrap Exactive and Orbitrap Exploris instruments were considered to ensure comparable acquisition characteristics. Datasets were required to contain Thermo RAW files with both MS1 and MS2 scans and to have valid GNPS dataset indicators. In total, 72 datasets meeting these inclusion criteria were retained, spanning 64 independent laboratories and encompassing diverse biological and environmental sample types (Supplementary Table S1).

Datasets were split at the laboratory level to avoid information leakage across splits. The training set consisted of 42 datasets from distinct laboratories, with runs randomly sampled and all associated MS2 scans included up to a maximum of 300,000 MS2 scans per dataset, amounting to a total number of 8,323,130 MS2 scans. The validation set consisted of 11 datasets from laboratories not represented in the training set with up to 100,000 MS2 scans per dataset, amounting to a total number of 905,432 MS2 scans. Three independent test sets (Test Set 1, 2, and 3) were constructed from additional unseen laboratories, consisting of 7, 5, and 7 datasets and 625,077, 416,751, and 677,205 MS2 scans, respectively.

RAW files were converted to the mzML format [18] using ThermoRawFileParser (version 1.4.5) [15]. Native peak picking was applied, and exception data (such as internal calibrant masses) were excluded.

MS1 spectra were extracted from the mzML files and preprocessed by filtering ion intensities below 1% of the base peak intensity, optionally truncating to a maximum of the 400 most intense ions, and scaling ion intensities using square root transformation. MS1 preprocessing was implemented using the spectrum_utils library (version 0.3.2) [4]. Precursor *m/z* values and charge states were extracted from the MS2 scan metadata for each MS2 scan following an MS1 scan.

Instrument configuration parameters were extracted directly from Thermo RAW files using ScanHeadsman [1]. For each RAW file, ScanHeadsman generated a per-scan CSV file containing acquisition and instrument metadata. A predefined subset of instrument parameters known to influence MS2 quality was retained for model training, including polarity, ionization mode, MS2 isolation width, ion injection time, Orbitrap resolution, AGC target, normalized HCD collision energy, precursor charge state, activation type, and related calibration parameters (Supplementary Table S2). Numerical instrument parameters were standardized using a global mean and standard deviation computed on the training set only. Categorical parameters were encoded using one-hot encoding. Scans with missing numerical or categorical instrument metadata were discarded.

### Data labeling and positive-unlabeled setting

MS2 scans were labeled using spectral library matching against the GNPS community spectral libraries (downloaded on 30/09/2025 and containing 880,127 reference library spectra), implemented using matchms (version 0.30.2) [14]. Library matching was performed using cosine similarity, precursor mass tolerance 0.05 Da, and fragment mass tolerance 0.05 Da. Scans with cosine similarity greater than 0.7 and at least six matched fragment ions were assigned positive labels (*Y* = 1), with all remaining scans assigned unlabeled status (*Y* = 0). Unlabeled scans include a mixture of low-quality spectra and high-quality spectra corresponding to compounds absent from the reference library.

This labeling scheme defines a PU learning problem, where only positive examples are reliably observed [3]. Let *X* ∈ ℝ^*d*^ denote the model input and *Y* ∈ {0, 1} the class label. Let *f* : ℝ^*d*^ → ℝ be an arbitrary real-valued decision function. Let *n*_*p*_ denote the number of positive samples and *n*_*u*_ denote the number of unlabeled samples in the training set. We define two subsets:

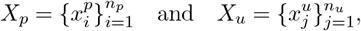

and the classifier is trained using both *X*_*p*_ (set of positive samples) and *X*_*u*_ (unlabeled samples).

The class-prior probability is defined as *π*_*p*_ = *P* (*Y* = 1). Using a binary cross-entropy loss ℓ(·), the positive and negative risks are defined as

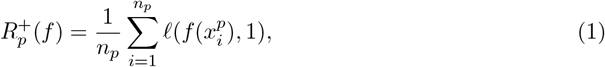

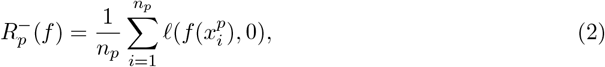

and

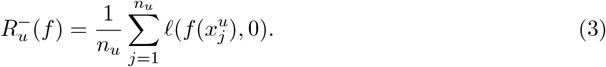

The nnPU risk [16] is defined as

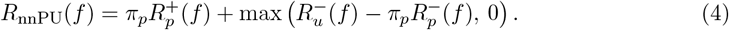

During training, if the estimated negative risk violated the non-negativity constraint by more than a threshold *β*, the optimization objective was modified to minimize the negative risk directly. The hyperparameter *β* was set to 0.

The class-prior probability *π*_*p*_ was estimated using the method of Elkan and Noto [11]. A preliminary classifier was trained to distinguish labeled positives from unlabeled samples using binary cross-entropy loss. The classifier was evaluated on known positive samples from Test Set 1 to estimate

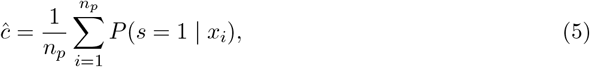

where *s* = 1 indicates a labeled sample. The class prior was then estimated as

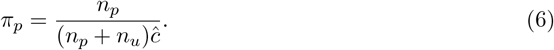

Class priors were estimated separately for positive and negative ionization modes.

### Model architecture and training

The model consists of three components: an MS1 spectrum encoder, an instrument settings encoder, and a feature integration and classification head (Figure 1b). Hyperparameters governing the architecture and training process were selected through a manual search across 12 distinct configurations. The search explored variations in the transformer embedding size, instrument setting embedding dimension, number of transformer encoder layers, learning rates, and optimizer choice. Each configuration was trained for a maximum of 30 epochs, with early stopping applied if the validation loss failed to improve. All configurations were evaluated on the validation set, and the final model was selected based on the minimum validation loss.

MS1 spectra were processed using a spectrum transformer-based encoder [23]. Each ion was embedded using its *m/z* value and normalized intensity. The precursor *m/z* was encoded as a global token and prepended to the sequence of peak embeddings. The transformer encoder comprised two layers, with 8 attention heads per layer and an embedding dimension of 128. Dropout with a rate of 0.3 was applied within the spectrum transformer encoder layers, between the layers of the instrument settings encoder, and to the combined representation before the final output layer. Instrument parameters were processed in parallel using a two-layer fully connected neural network with hidden dimension 16 in both layers and ReLU activations. The final MS1 embedding and instrument embedding were concatenated and passed through an integration layer with ReLU activation. A final linear layer produced a single logit, which was transformed into a probability using a sigmoid function. The final architecture corresponds to the best-performing configuration identified during hyperparameter optimization.

The model was trained using the AdamW optimizer with a learning rate of 5.0 *×* 10^−6^ for the transformer encoder and 1.0 *×* 10^−5^ for the linear layers, with a weight decay of 0.001 and batch size 256. Each batch contained a mixture of positive and unlabeled samples. Training was performed for 30 epochs with early stopping based on validation recall. All experiments were conducted on a high-performance computing cluster node equipped with dual Intel Xeon Gold 5320 CPUs (2.20 GHz) and NVIDIA A100 GPUs (80 GB). Random seeds were fixed to ensure reproducibility.

A decision threshold on predicted probabilities was selected using Test Set 2. The threshold was defined as the 5th percentile of predicted probabilities among known positive samples in Test Set 2. This strategy prioritizes high recall for positive spectra while remaining independent of the final benchmark evaluation.

### Evaluation and validation

Model performance was assessed on Test Set 3 by measuring recall on known positive samples using the decision threshold defined on Test Set 2. Because true negative labels are unavailable, additional validation was performed by examining independent indicators of MS2 spectral quality on the unlabeled portion of Test Set 3.

Structural explainability was evaluated using SIRIUS (version 6.3.3) [9]. Unless otherwise specified, default parameters were applied using the “Orbitrap” algorithm profile on input spectra converted to MGF format. Notably, the precursor mass tolerance for the identity search was set to 10.0 ppm (stricter than the default 20.0 ppm), while the fragment mass tolerance was set to 5.0 ppm. Spectral, formula, and structure searches were performed against the SIRIUS biomolecule structure database (bioDB). As a proxy for spectral interpretability, we extracted the “explained intensity” reported by SIRIUS, which quantifies the proportion of the MS2 total ion current (TIC) that can be explained by the inferred molecular formula. Only spectra for which SIRIUS returned at least one candidate formula were included in this analysis.

Fragment ion richness was assessed by counting the number of fragment ions with non-zero intensity in each MS2 spectrum. No additional intensity thresholding was applied beyond the preprocessing already performed during mzML conversion. Spectra containing fewer than three non-zero intensity fragment ions were considered to exhibit low fragment ion content.

Precursor charge state was obtained directly from the MS2 scan metadata in the mzML files when available. Only scans for which the instrument-assigned precursor charge state was present were included in this analysis. Spectra with an absolute precursor charge state greater than two (|*z*|*>* 2) were flagged as likely non-metabolite contaminants. While such spectra can be of high quality, they typically correspond to multiply charged peptides or macromolecules, which fall outside the scope of metabolomics.

Spectral interference was estimated using the TIC percentage attributed to fragment ions occurring above the precursor *m/z* minus 5 Da. This region is not expected to contain true fragment ions under typical collision-induced dissociation or higher-energy collisional dissociation conditions and therefore serves as a proxy of chimeric fragmentation or co-isolation interference. Only singly charged precursor ions with non-zero interference TIC were included in this analysis to reduce confounding effects from multiply charged precursors.

Associations between model-predicted probabilities and quality indicators were evaluated using Pearson correlation. Statistical significance was assessed at a significance level of *α* = 0.05.

### Software and data availability

All code used in this study is available as open source under the Apache 2.0 license at https://github.com/bittremieuxlab/pu_ms2_identifiability. Processed datasets and trained model weights are available on Zenodo at https://doi.org/10.5281/zenodo.18266932.

## 3 Results

### Public metabolomics data exhibits low spectrum annotation rates

The final training, validation, and test splits comprised 10,947,595 MS2 scans collected across dozens of independent laboratories, reflecting a wide range of sample types and acquisition conditions (Supplementary Table S1). Across all splits, only a minority of scans were assigned positive labels through spectral library matching, while the majority remained unlabeled (Figure 2). This pronounced imbalance underscores the difficulty of the task and motivates the use of PU learning rather than fully supervised classification. Importantly, all evaluations reported below were performed on datasets originating from laboratories that were not represented during training, ensuring a stringent assessment of model generalization.

**Figure 2.**
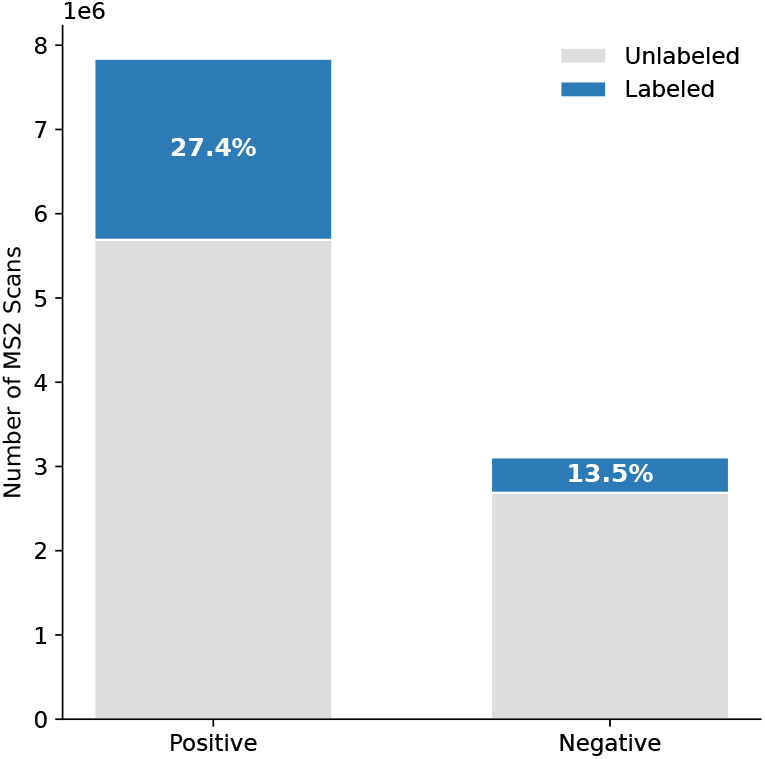
Total count of MS2 scans split by ionization polarity (Positive vs. Negative). Colors distinguish between scans with ground-truth labels derived from spectral library matching (blue) and those that remain unlabeled (grey).

### Ionization mode-dependent expected fraction of identifiable MS2 spectra

A key component of nnPU learning is the estimation of the class-prior probability *π*_*p*_. Following the approach of Elkan and Noto [11], preliminary classifiers were trained to discriminate labeled positive scans from unlabeled scans using binary cross-entropy loss. These classifiers were evaluated on known positive samples from the independent Test Set 1 to estimate the probability *c* = *P* (*s* = 1 | *y* = 1), corresponding to the probability that a truly positive spectrum is labeled.

Separate estimates were obtained for positive and negative ionization modes. For positive ionization mode, the estimated value was *ĉ* = 0.5889, resulting in a class-prior estimate of *π*_*p*_ = 0.454, whereas for negative ionization mode *ĉ* = 0.4486, corresponding to *π*_*p*_ = 0.285. These differences reflect well-known polarity-dependent differences in accessible chemical space, fragmentation efficiency, and reference library representation [17], all of which influence the likelihood that an MS2 spectrum is both informative and successfully identified, thereby motivating the use of polarity-specific class priors during nnPU training.

### nnPU learning recovers identifiable spectra in unseen laboratories

To define a probability threshold for downstream analyses without biasing the final evaluation, threshold selection was performed using an independent dataset (Test Set 2). The threshold was set to the 5th percentile of predicted probabilities among known positive samples in this dataset, yielding a value of 0.767 (Figure 3a). This operating point was chosen to prioritize high recall of identifiable spectra, consistent with the intended use of the model as a tool for assessing and optimizing MS2 acquisition quality rather than aggressively filtering data.

**Figure 3.**
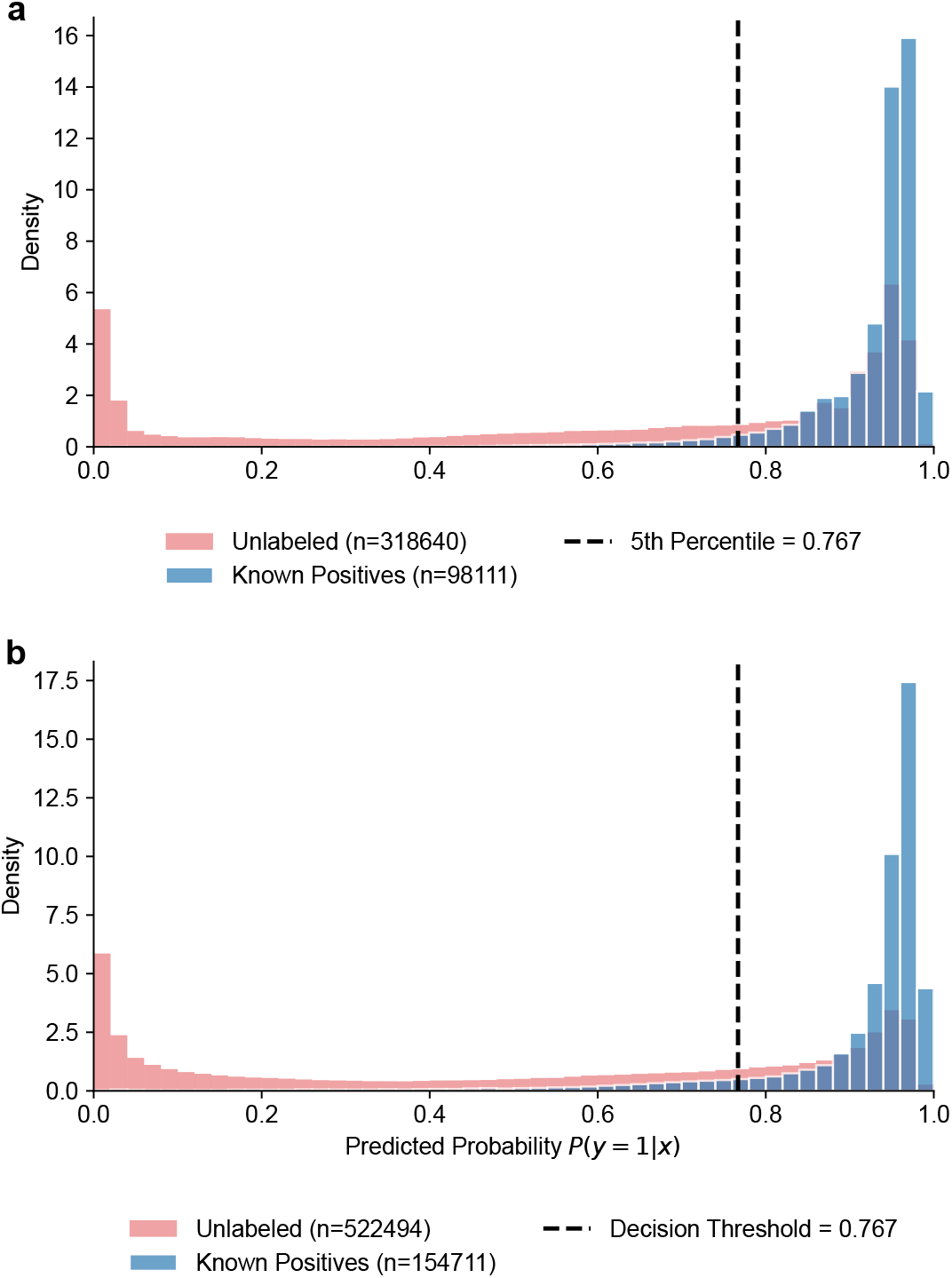
**(a)** Distribution of predicted probabilities for known positive and unlabeled spectra in Test Set 2, used to define the decision threshold without biasing final evaluation. **(b)** Distribution of predicted probabilities for known positive and unlabeled spectra in the held-out Test Set 3, demonstrating effective retrieval of known high-quality spectra and stratification of potentially high-quality spectra within the unlabeled set under previously unseen experimental conditions.

Model performance was subsequently evaluated on Test Set 3, which served as a fully heldout benchmark comprising datasets from previously unseen laboratories. Using the predefined decision threshold, the model achieved a recall of 0.8986 on known positive spectra, reflecting comparable performance across independent datasets (Figure 3b). The distribution of predicted probabilities exhibited a clear bimodal structure, with the majority of labeled positive spectra assigned high probabilities and successfully retrieved above the decision threshold. These results demonstrate that the model generalizes across laboratories and acquisition conditions and successfully identifies the majority of spectra that are known to be of high quality.

In addition to correctly identifying known positives, the model assigned high probabilities to a subset of unlabeled scans (approximately 37.61%). These spectra represent putative “hidden positives,” that is, high-quality spectra corresponding to molecular species absent from the reference library. This behavior is expected under the PU learning framework and indicates that the model does not trivially equate the absence of a library match with low spectral quality.

### Quality prediction corresponds to structural explainability of MS2 spectra

To assess whether model predictions align with chemically meaningful notions of spectral quality beyond library matching, we evaluated structural explainability using SIRIUS [9]. While SIRIUS addresses a conceptually similar task to spectral library searching, namely the annotation of MS2 spectra, it operates through a fundamentally different mechanism. Rather than relying on reference spectral libraries, SIRIUS performs molecular formula inference by searching against large structural databases and evaluating the consistency of observed fragment ions with plausible fragmentation trees. Because structural databases are substantially broader in coverage than reference spectral libraries, SIRIUS is expected to successfully assign molecular formulas to a wider range of chemically valid, high-quality spectra, including many that remain unmatched in library-based searches. Consequently, we hypothesized that spectra assigned higher quality probabilities by our model would, on average, exhibit greater structural explainability as quantified by SIRIUS. At the same time, structural databases remain incomplete [19] and fragmentation models are imperfect, such that the absence of a SIRIUS assignment does not necessarily imply poor spectral quality, leading to a similar PU setting.

Across unlabeled scans in Test Set 3, the model-predicted probability of high quality exhibited a positive correlation with the explained intensity. Specifically, Pearson correlation analysis yielded a correlation coefficient of *r* = 0.28 with *p* ≪ 0.001 (Figure 4a). Spectra assigned higher probabilities by the model were therefore more likely to produce fragment-rich spectra amenable to structural interpretation. This result provides independent validation that the model captures chemically relevant aspects of MS2 quality rather than merely reproducing library matching behavior.

**Figure 4.**
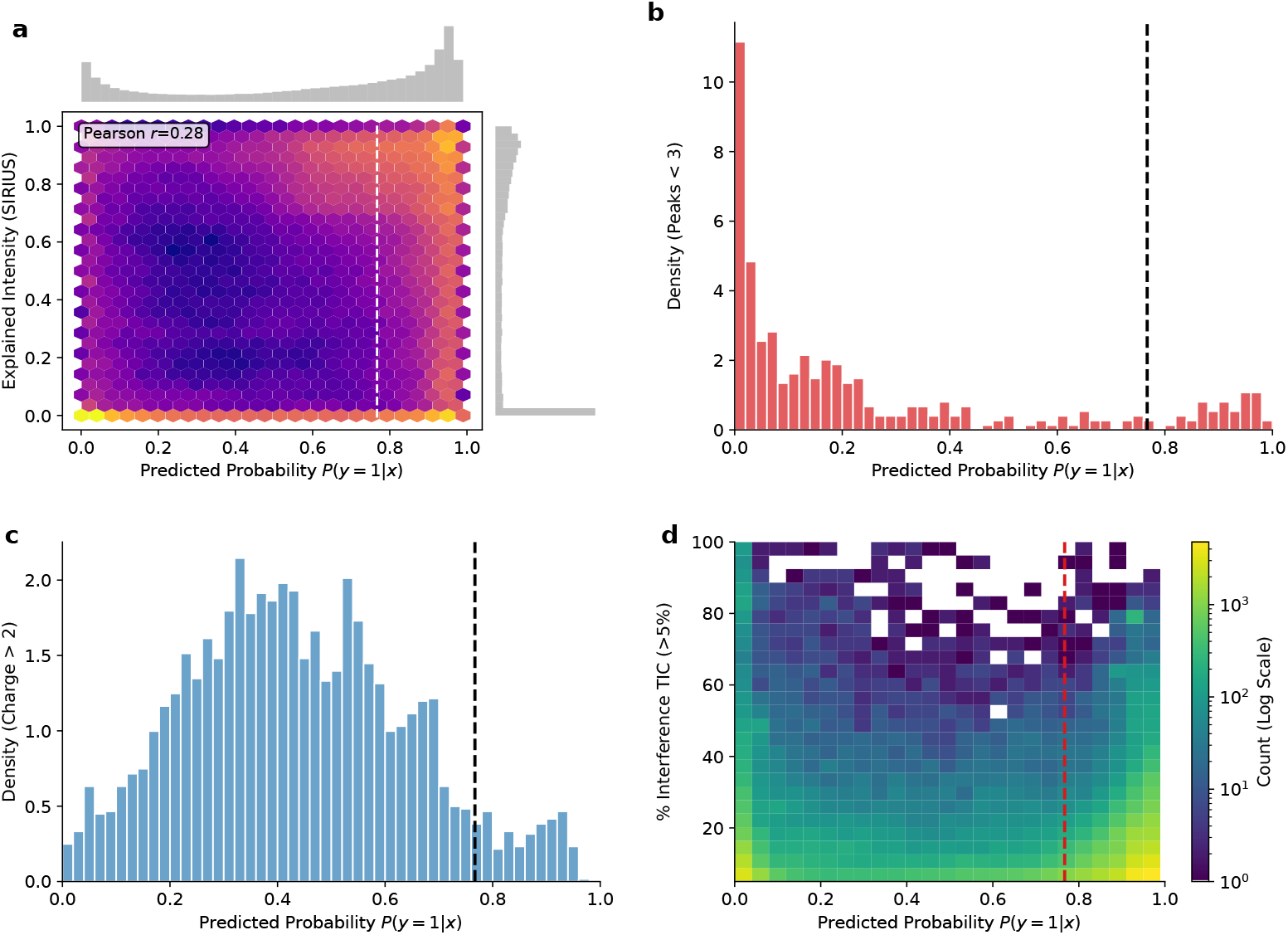
Model validation using independent indicators of MS2 spectral quality. **(a)** Joint density of SIRIUS structural explainability (explained intensity) and predicted probability for unlabeled MS2 scans. **(b)** Distribution of predicted probabilities for unlabeled MS2 scans exhibiting sparse fragmentation, defined as fewer than three non-zero intensity fragment ions. **(c)** Distribution of predicted probabilities for unlabeled MS2 scans with high precursor charge states. **(d)** Joint density of predicted probability and spectral interference, quantified as the percentage of total ion current observed in the window starting 5 Da below the precursor m/z, for singly charged unlabeled MS2 scans with with interference greater than 5%.

### Low predicted probabilities match MS2 quality heuristics

In addition to structural explainability, we examined several established MS2 spectrum-level quality heuristics to further characterize model predictions on the unlabeled set. These analyses focused on fragment ion richness, precursor charge state, and spectral interference, all of which are known to influence MS2 interpretability and are computed directly from the acquired MS2 spectra. Importantly, the predictive model itself did not have access to any MS2 information and relied exclusively on MS1 context and instrument parameters, such that agreement with these MS2-derived heuristics reflects the model’s ability to anticipate downstream spectral quality rather than post hoc evaluation of the fragmentation data.

First, fragment ion richness was assessed by counting the number of non-zero intensity fragment ions in each MS2 spectrum (Figure 4b). Spectra predicted with low probability were substantially more likely to contain very few fragment ions. Second, precursor charge state was examined for scans with instrument-assigned charge annotations (Figure 4c). High precursor charge states ( |*z*| *>* 2), which are often associated with poor fragmentation or ambiguous spectral interpretation in small molecule MS2 data, were markedly more frequent among spectra predicted to be of low quality. Finally, spectral interference was estimated using the TIC fraction arising from fragment ions above the precursor *m/z* minus 5 Da. Among singly charged precursors with non-zero interference, we observed a weak but statistically significant negative correlation between predicted probability and interference TIC (*r* = −0.0912, *p* ≪ 0.001) (Figure 4d). This indicates that increased estimated co-isolation or chimeric fragmentation tends to reduce predicted MS2 quality.

Taken together, these heuristic analyses demonstrate that spectra assigned low probability by the model exhibit multiple independent indicators of poor spectral quality, while high-probability spectra are enriched for physically and chemically plausible fragmentation patterns.

## 4 Discussion and conclusions

In this work, we introduced a learned approach to predict the likelihood that a tandem mass spectrum will be of sufficient quality to support confident metabolite identification, using only information available prior to MS2 interpretation. By combining MS1 spectral context with instrument acquisition parameters and formulating the task as a PU learning problem, we demonstrate that it is possible to estimate MS2 identifiability without relying on explicit negative labels or post hoc analysis of the MS2 spectrum itself. Across large-scale, laboratory-disjoint benchmark datasets, the proposed model achieved high recall on known positive spectra and exhibited consistent, interpretable behavior on unlabeled data.

A central contribution of this study is the explicit treatment of label uncertainty inherent to untargeted metabolomics. Unlike supervised settings where both positive and negative labels are well defined, metabolomics data contain only a sparse and biased set of confidently identified spectra, while the majority of scans remain unlabeled for reasons that are not directly tied to spectral quality. By adopting an nnPU learning framework, the model is able to leverage reliable positives while accounting for the mixed nature of the unlabeled pool. The use of polarity-specific class-prior estimates further reflects known differences between ionization modes and improves model calibration across heterogeneous datasets.

The results suggest that the model captures meaningful aspects of MS2 identifiability rather than artifacts of dataset composition or laboratory-specific acquisition practices. Predicted probabilities generalized well to data acquired in previously unseen laboratories, indicating that the learned representations reflect common relationships between precursor context, acquisition parameters, and downstream spectral informativeness. Importantly, validation using independent quality indicators provides supporting evidence for this interpretation. Spectra assigned low predicted probabilities were enriched for sparse fragmentation, atypical precursor charge states, and spectral interference, while spectra assigned higher probabilities were more likely to exhibit chemically interpretable fragmentation patterns as quantified by SIRIUS explainability metrics.

It is important to emphasize what the proposed model does and does not predict. The output probability should be interpreted as an estimate of MS2 identifiability under typical library-based and structure-inference workflows, rather than as an absolute or intrinsic measure of spectral quality. The model does not infer molecular structure, nor does it guarantee successful identification for any individual spectrum. Instead, it integrates precursor-level information and acquisition settings to estimate whether the resulting MS2 scan is likely to be informative, recognizing that MS2 quality is a multifactorial and probabilistic property. The moderate strength of correlations with individual quality indicators is therefore expected and reflects the complexity of the underlying phenomenon rather than a limitation of the approach.

From a practical perspective, this work highlights the potential of predictive models to inform mass spectrometry acquisition strategies. Because the model operates on MS1 spectra and instrument parameters alone, it can be applied prior to or independently of MS2 interpretation. This opens the possibility of computationally exploring acquisition parameter spaces, prioritizing MS2 scans that are more likely to yield interpretable spectra, or guiding adaptive acquisition strategies that respond to precursor context in real time [24]. Such applications are complementary to experimental optimization and post hoc quality control, offering a means to reduce the acquisition of uninformative spectra and improve overall annotation efficiency.

Several limitations of the present study should be acknowledged. First, positive labels are derived from spectral library matches and are therefore biased toward compound classes and acquisition conditions that are well represented in existing libraries. The class-prior estimation relies on the “selected completely at random” assumption, which may be violated in practice if labeling probability depends on spectral or chemical properties not fully captured by the model inputs. However, prior theoretical and empirical work on nnPU learning indicates that its optimization is relatively robust to moderate misspecification of the class-prior probability, particularly when the learning objective is dominated by the positive risk and the non-negativity constraint prevents overfitting of the negative risk [16]. Second, the analysis is restricted to Thermo Fisher Scientific Orbitrap Exactive and Exploris instruments, and model behavior on other platforms or fragmentation modalities remains to be evaluated. Moreover, public datasets likely reflect common acquisition practices rather than the full parameter space; targeted sampling of underrepresented parameter combinations, either offline or through adaptive online acquisition, could improve model robustness.

Future work will focus on extending and validating this framework along two closely connected directions. First, prospective acquisition studies will be required to determine whether instrument parameter choices informed by model predictions lead to measurable improvements in MS2 annotation rates and overall spectral interpretability under real experimental conditions. Such studies are essential to establish the practical impact of predictive quality assessment beyond retrospective analysis. Second, successful validation in controlled acquisition experiments would naturally motivate the integration of predictive MS2 quality estimation into real-time or iterative acquisition workflows. In this setting, predicted quality scores could be used to dynamically guide fragmentation decisions or acquisition parameter adjustments, enabling data-driven optimization of MS2 acquisition during ongoing experiments.

In conclusion, this study demonstrates that MS2 identifiability in untargeted metabolomics can be meaningfully predicted from MS1 context and acquisition parameters alone, despite the absence of explicit negative labels. By combining large-scale public data with a principled learning framework, we provide evidence that MS2 spectral quality can be meaningfully predicted from precursor characteristics and acquisition parameters alone, prior to inspecting the fragmentation spectrum itself. We anticipate that this perspective will contribute to the development of model-informed data acquisition strategies and help bridge the gap between experimental design and computational analysis in metabolomics.

## 5 Competing interests

No competing interest is declared.

## 6 Author contributions statement

M.B.: data curation, investigation, methodology, software, validation, visualization, writing – original draft, writing – review & editing. T.J.: methodology, software, writing – review & editing. L.F.N.: conceptualization, funding acquisition, methodology, project administration, resources, supervision, writing – review & editing. W.B.: conceptualization, funding acquisition, methodology, project administration, resources, supervision, writing – original draft, writing – review & editing.

## 7 Acknowledgments

M.B. was supported by a PhD fellowship from the Interdisciplinary Institute for Artificial Intelligence (3IA) Côte d’Azur (ANR-19-P3IA-0002 and ANR-23-IACL-0001) and the Initiative of Excellence Université Côte d’Azur (ANR-15-IDEX-01). This work is supported in part by funds from the Research Foundation–Flanders (FWO G0AHY25N). This work was supported in part by the French government through the France 2030 investment plan managed by the National Research Agency (ANR) under reference ANR-22-CPJ2-0048-01. The authors are grateful to the Université Côte d’Azur’s Center for High-Performance Computing (OPAL infrastructure) for providing resources and support.

## Notes

### Competing Interest Statement

The authors have declared no competing interest.

