## Supplementary Table S1, Supplementary Table S2 for "Positive-Unlabeled Learning for Predicting Small Molecule MS2 Identifiability from MS1 Context and Acquisition Parameters"

### Supplementary Notes

| ID | Lab / PI | Species | mzMLs | MS2 Scans (Used/Total) |
|---|---|---|---|---|
| MSV000096347 | Paul Abraham;ORNL;United States | Populus trichocarpa (NCBITaxon:3694) | 38 | 299565/7803163 |
| MSV000098437 | Shirley M. Tsunoda;University of California San Diego;USA | Homo sapiens (NCBITaxon:9606) | 101 | 299990/4105109 |
| MSV000098293 | Pam Ronald;UC Davis;United States | Arabidopsis | 75 | 299903/3752184 |
| MSV000096860 | Weisi;Nanjing University;China | environmental sample wastewater | 18 | 299844/3589760 |
| MSV000098287 | Gabriel Castrillo | Spirodela polyrhiza | 37 | 298344/2486971 |
| MSV000096359 | Lindsey Burnett;University of California San Diego;United States | Homo sapiens (NCBITaxon:9606) | 94 | 297408/2374064 |
| MSV000098286 | Debashish Bhattacharya | soil microbiome | 50 | 298861/1837462 |
| MSV000096317 | Alicia Purcell;Northern Arizona University;United States | glacier microbiome | 50 | 297401/1504582 |
| MSV000092256 | Dean E. Riechers | Triticum aestivum (NCBITaxon:4565) | 20 | 286399/1360883 |
| MSV000099130 | Mauricio Caraballo | Several | 53 | 296325/897361 |
| MSV000098289 | Tiffany Lowe-Power;UC Davis;United States | Solanum lycopersicum root exudate | 44 | 297837/879127 |
| MSV000088843 | Fabian Rivera | Mus sp. (NCBITaxon:10095) | 134 | 287582/364110 |
| MSV000096729 | David Gonzalez;UCSD;United States | Homo sapiens (NCBITaxon:9606) | 105 | 281285/281285 |
| MSV000088723 | Aaron Hartmann | environmental samples <Bacillariophyta> (NCBITaxon:33858) | 40 | 234156/267476 |
| MSV000098518 | Mi Zhang;Toho University;Japan\|Kouharu Otsuki;Toho University;Japan\|Takashi Kikuchi;Toho University;Japan\|Wei Li;Toho University;Japan | Daphne (NCBITaxon:66679)Wikstroemia (NCBITaxon:142693)Stellera (NCBITaxon:142737)Edgeworthia (NCBITaxon:142180) | 124 | 239862/239862 |
| MSV000086956 | Hosein Mohimani | 296\|Pseudomonas fragi | 36 | 27117/27117 |
| MSV000098306 | Edward T. Chouchani;Dana Farber Cancer Institue;USA | Mus musculus (NCBITaxon:10090)Homo sapiens (NCBITaxon:9606) | 2 | 19225/19225 |
| MSV000097074 | Mariana Alves Reis;CIIMAR;Portugal | Lusitaniella coriacea LEGE 07167 | 5 | 12274/12274 |
| MSV000096582 | Xiaoyang Su;Rutgers Cancer Institute of New Jersey;USA | Montipora capitata (NCBITaxon:46704)Pocillopora acuta (NCBITaxon:1491507) | 3 | 24668/24668 |
| MSV000093813 | A.J. Marian Walhout | Caenorhabditis elegans (NCBITaxon:6239) | 8 | 94864/94864 |

*Continued on next page...*

| ID | Lab / PI | Species | mzMLs | MS2 Scans (Used/Total) |
|---|---|---|---|---|
| MSV000083766 | Bernhard Palsson | Staphylococcus aureus (NCBITaxon:1280) | 52 | 110250/159622 |
| MSV000092254 | Dr. Edward Owusu-Ansah | Drosophila melanogaster (NCBITaxon:7227) | 6 | 13705/13705 |
| MSV000083372 | Forest Rohwer | Coral | 68 | 148706/154102 |
| MSV000087702 | Georg Pohnert | Synechococcus (NCBITaxon:1129)environmental samples <Bacillariophyta> (NCBITaxon:33858) | 36 | 169083/193762 |
| MSV000088937 | Daniel Petras | environmental samples <Apicomplexa> (NCBITaxon:282487) | 59 | 298787/1759211 |
| MSV000088732 | Daniel Petras | unknown species | 58 | 299921/559185 |
| MSV000098291 | Sabeeha Merchant;UC Berkeley;United States | Chlamydomonas reinhardtii CC5390 Brevundimonas sp. | 16 | 98994/3232698 |
| MSV000097708 | Sarkis Mazmanian;California Institute of Technology;United States | Mus musculus (NCBITaxon:10090) | 24 | 99791/1406247 |
| MSV000084364 | Jacob Agerbo Rasmussen | Oncorhynchus mykiss (NCBITaxon:8022) | 7 | 95180/1141177 |
| MSV000098284 | Jonelle Basso;Lawrence Berkeley National Laboratory;United States | Arabidopsis microbiome | 19 | 98928/896645 |
| MSV000090950 | Amir Zarrinpar;University of California San Diego;USA | Mus musculus (NCBITaxon:10090) | 60 | 99831/386700 |
| MSV000098817 | Hiroshi Otani;LBL;United States | Streptomyces coelicolor | 19 | 99795/252923 |
| MSV000099114 | Allegra Aron | Methylobacterium aquaticum (NCBITaxon:270351) | 7 | 26633/37869 |
| MSV000095703 | Bret Cooper | Penicillium (NCBITaxon:5073) | 10 | 81719/81719 |
| MSV000095217 | Zachary A. Quinlan;University of Hawai'i at Manoa;United States | Montipora capitata (NCBITaxon:46704)Porites lobata (NCBITaxon:104759) | 45 | 99883/2073990 |
| MSV000094782 | Esther Singer;Lawrence Berkeley National Laboratory;United States | Panicum hallii microbiome | 25 | 98600/631246 |
| MSV000095029 | Zachary Quinlan;Hawaii Institute of Marine Biology;USA | Scleractinia (NCBITaxon:6125)Montipora (NCBITaxon:46703)Porites (NCBITaxon:46719) | 45 | 99812/2024429 |
| MSV000095207 | Jorge Barriuso;CIB Center for Biological Research;Spain | Ophiostoma piceae CECT 20146 Pseudomonas putida KT2440 | 17 | 98210/1572431 |
| MSV000094929 | Sibgha Tayyab;Tubingen University;Germany | Staphylococcus aureus (NCBITaxon:1280) | 19 | 99856/184505 |
| MSV000097243 | Lihini Aluwihare | environmental samples <Bacillariophyta> (NCBITaxon:33858) | 21 | 99483/1341939 |

*Continued on next page...*

| ID | Lab / PI | Species | mzMLs | MS2 Scans (Used/Total) |
|---|---|---|---|---|
| MSV000094923 | sibgha sibgha;Tubingen University;Germany | Escherichia coli (NCBITaxon:562) | 4 | 24209/28980 |
| MSV000095461 | Andrew Truman;John Innes Centre;United Kingdom | Pseudomonas sp. Ps652 | 35 | 97903/174357 |
| MSV000096694 | Karen Pierce | Homo sapiens (NCBITaxon:9606) | 18 | 96253/605665 |
| MSV000094781 | Davinia Salvachua;National Renewable Energy Laboratory;United States | Trametes versicolor Ceriporiopsis subvermispora | 23 | 99994/2464577 |
| MSV000094780 | Kristen DeAngelis;University of Massachusets;United States | soil incubated with Actinobacteria (specific genuses: Kitasatospora Streptomyces Leifsonia sp. BS71 Streptacidiphilus Corynebacteria Planotetraspora Frankia Catenulispora) | 17 | 98909/3256571 |
| MSV000094763 | Alyson Santoro;University of California Santa Barbara;United States | Nitrosopumilus adriaticus CCS1 Nitrospina gracilis Nb-211 | 18 | 99173/1945579 |
| MSV000094574 | Benjamin C Orsburn;Johns Hopkins;USA | Homo sapiens (NCBITaxon:9606) | 42 | 86601/86601 |
| MSV000094118 | Jo Handelsman;UW-Madison;US\|Marc Chevrette;University of Florida;United States | Pseudomonas koreensis (NCBITaxon:198620)Bacillus cereus (NCBITaxon:1396)Flavobacterium johnsoniae (NCBITaxon:986) | 13 | 96088/564390 |
| MSV000094091 | Philipp Zerbe;UC Davis;United States | Panicum virgatum | 25 | 97026/805443 |
| MSV000094089 | Aymerick Eudes;JBEI;United States | Populus | 17 | 99414/1695514 |

**Table S1:** The metadata for the datasets used for training and testing. This includes the Principal Investigator (PI) who shared the dataset, the species, the number of mzML files considered from that dataset, and the number of MS2 scans considered by the model out of the total available scans MS2.

| Parameter | Description |
|---|---|
| Polarity | Indicates the polarity of the ion detection, either positive or negative. |
| Ionization | Type of ion source used for ion generation, including ESI or NSI. |
| MS2 Isolation Width | The $m/z$ window applied for precursor ion selection. |
| Ion Injection Time (ms) | Maximum injection time allowed for ion accumulation in the C-trap. |
| Conversion Parameter C | Calibration-related parameter for signal conversion. |
| Mild Trapping Mode | Indicates whether mild trapping mode was enabled during ion accumulation. |
| Orbitrap Resolution | Resolving power of the Orbitrap analyzer at a defined $m/z$. |
| AGC Target | Automatic gain control (AGC) represents the number of ions filling the C-trap before being transferred into the Orbitrap. |
| Normalized HCD Energy | Percentage of energy applied for precursor fragmentation in the collision cell. |
| LM $m/z$-Correction (ppm) | Correction factor applied to low-mass ions, expressed in parts per million (ppm). |
| Activation 1 | The method used to fragment ions in MS/MS analysis, such as CID or HCD. |

**Table S2:** Instrument parameter settings and descriptions.
